# Identification of the maize Mediator CDK8 module, and *Dissociation* insertional mutagenesis of *ZmMed12a*

**DOI:** 10.1101/097204

**Authors:** Tania Núñez-Ríos, Kevin R. Ahern, Ana Laura Alonso-Nieves, Daniel Lepe-Soltero, Carol Martínez-Camacho, Marcelina García-Aguilar, Thomas P. Brutnell, C. Stewart Gillmor, Ruairidh James Hay Sawers

## Abstract

Mediator is a conserved transcriptional co-activator that links transcription factors bound at enhancer elements to RNA Polymerase II. Mediator-RNA Polymerase II interactions can be sterically hindered by the Cyclin Dependent Kinase 8 (CDK8) module, a submodule of Mediator that acts to repress transcription in response to discrete cellular and environmental cues. The CDK8 module is conserved in all eukaryotes and consists of 4 proteins: CDK8, CYCLIN C (CYCC), MED12, and MED13. In this study, we have characterized the CDK8 module of Mediator in maize. The maize genome contains single copy genes for *Cdk8, CycC,* and *Med13,* and two genes for *Med12.* Analysis of expression data for the CDK8 module demonstrated that all five genes are broadly expressed in maize tissues, with *ZmMed12a, ZmMed12b,* and *ZmMed13* exhibiting similar expression patterns. We performed a *Dissociation (Ds)* insertional mutagenesis, recovering two independent insertions in the *ZmMed12a* gene. One of these *Ds* insertions results in a truncation of the *ZmMed12a* transcript. Our molecular characterization of the maize CDK8 module, as well as transposon tagging of *ZmMed12a,* establish the basis for molecular and functional studies of these important transcriptional regulators in *Zea mays*.

## INTRODUCTION

Transcriptional regulation plays an essential role in almost all aspects of development and physiology, including responses to the biotic and abiotic environment. One key regulator of transcription is Mediator, a multiprotein complex conserved from yeast to plants to animals, which was initially identified based on its requirement for transcription of virtually all protein-coding genes (Kelleher et al., 1990; Flanagan et al., 1991; Bourbon, 2008). The Core Mediator consists of Head, Middle and Tail domains, and typically functions as a transcriptional co-activator, linking transcription factors bound at upstream enhancer elements to RNA polymerase II (RNA pol II) (reviewed in Yin and Wang, 2014; Allen and Taatjes, 2015). The Head and Middle domains interact with RNA pol II, while the Tail domain is thought to interact with specific transcription factors (Tsai et al., 2014; Robinson et al., 2015; Plaschka et al., 2015; reviewed in Larivière et al., 2012). A fourth Mediator module shows transient association with Core Mediator and often acts to repress transcription. This Cyclin Dependent Kinase 8 (CDK8) module is composed of the proteins MED12, MED13, CYCLIN C (CYCC), and CDK8 (reviewed in Björklund and Gustafsson, 2005). In agreement with the variable association of the CDK8 module with Core Mediator, purification of Mediator from *Arabidopsis thaliana* yielded both conserved Core Mediator subunits, as well as subunits unique to Arabidopsis, but did not include components of the CDK8 module (Bäckström et al., 2007).

In yeast and animals, components of the CDK8 module can regulate transcription in several ways, with different subunits playing different roles. One mechanism for transcriptional repression involves steric inhibition, where the CDK8 module occupies the Core Mediator pocket that binds RNA pol II, thereby preventing interaction of Core Mediator and RNA pol II (Elmlund et al., 2006; Tsai et al., 2013). Transcriptional repression by this steric mechanism has the potential to be dynamic, as the occupancy of the RNA pol II binding pocket can be modulated during subsequent rounds of assembly of the Mediator-RNA pol II holoenzyme (reviewed in Allen and Taatjes, 2015). This steric mechanism involves all four units of the CDK8 module, with the MED13 subunit playing the most important role, interacting directly with the Middle domain of Core Mediator (Knuesel et al., 2009; Tsai et al., 2013). The MED13 subunit also serves an important function in regulation of CDK8 module stability: phosphorylation of a conserved phosphodegron site in MED13 can lead to recognition by a ubiquitin ligase complex, and subsequent degradation (Davis et al., 2013).

In Arabidopsis, components of the CDK8 module were initially identified by their requirement for development, and also affect the response to fungal pathogens and cellular stress. Mutations in *CDK8* were identified as enhancers of the phenotype of the floral homeotic mutant *hua1hua2,* and thus were named *hua enhancer 3 (hen3)*. *hen3* mutants affect floral organ identity, as well as leaf size and cell shape, and the HEN3 protein was demonstrated to have CDK8 kinase activity (Wang and Chen, 2004). CDK8 regulates retrograde signaling from the mitochondria to the nucleus in response to H_2_O_2_ and cold stress (Ng et al., 2013). *CDK8*, as well as *MED12* and *MED13*, are also required for the response to both fungal and bacterial pathogens (Zhu et al., 2014).

Mutations in *MED12* and *MED13* were initially reported from a genetic screen for regulators of pattern formation in Arabidopsis embryogenesis, and were named *center city (cct)* and *grand central (gct)*, to reflect the increased size of the shoot apical meristem (SAM) in these mutants. *cct* and *gct* mutants delay the timing of pattern formation during embryogenesis, rather than affecting pattern formation *per se-* the increased size of the SAM in *cct* and *gct* mutants can be attributed to its formation later in embryogenesis compared to the wild type (wt) (Gillmor et al., 2010). The delayed formation of the SAM may be related to auxin signaling, as both the *med13* allele *macchi-bou2 (mab2)*, and the *med12* allele *cryptic precocious* (*crp*) act as enhancers of a mutation in the auxin dependent kinase *PINOID* (Furutani et al., 2004; Ito et al., 2011; Imura et al., 2012). Importantly for mechanistic studies of CDK8 module function in Arabidopsis, Ito et al. (2011) demonstrated that the MED13 and CDK8 proteins are both able to interact with Cyclin C, as has previously been demonstrated in Drosophila (Loncle et al., 2007). Consistent with studies showing auxin-related phenotypes for mutants in *MED12* and *MED13*, a recent study showed that both of these genes, as well as *CDK8*, are involved in auxin transcriptional responses, and that the MED13 protein relays signals from the IAA14 protein to repress the auxin responsive transcription factors ARF7 and ARF19 (Ito et al., 2016).

In addition to affecting the timing of pattern formation in embryogenesis, *MED12* and *MED13* also regulate the timing of post-embryonic phase transitions in Arabidopsis. A dominant allele of *med12* (named *cryptic precocious* (*crp-1D*)) was isolated in a genetic screen for enhancers of the early flowering phenotype conditioned by overexpression of the florigen *FT* (Imura et al., 2012). Loss of function mutants in *crp/cct* and *gct* show late flowering due to overexpression of the floral repressor *FLOWERING LOCUS C (FLC)*, as well as decreased expression of the floral promoters *FLOWERING LOCUS (FT)*, *TWIN SISTER OF FT (TSF)*, *SUPPRESSOR OF OVEREXPRESSION OF CONSTANTS 1 (SOC1)*, *APETALA 1 (AP1)* and *FRUITFULL (FUL)* (Imura et al., 2012; Gillmor et al., 2014). *cct* and *gct* mutants also misexpress seed specific genes during seedling development, and have an elongated vegetative phase due to overexpression of the microRNA miR156 (Gillmor et al., 2014), a master regulator of the vegetative phase in plants (Wu et al., 2009). Taken together, these results demonstrate that *MED12* and *MED13* act as master regulators of developmental timing in plants, regulating the timing of pattern formation in embryogenesis, the seed-to-seedling transition, vegetative phase change, and the transition to flowering (Gillmor et al., 2010; Ito et al., 2011; Imura et al., 2012; Gillmor et al., 2014).

Due to its importance in plant development and physiology, we have extended studies of the CDK8 module to the crop plant maize (*Zea mays*). Establishment of molecular and genetic resources for the study of the maize CDK8 module will allow evaluation of its role in the regulation of agricultural traits such as timing of flowering and seed development, as well as responses to biotic and abiotic stresses. One of the primary goals of this work was isolation of loss of function mutant alleles of maize CDK8 module-encoding genes. In maize, resources based on endogenous DNA transposons constitute the most accessible and widely-used technology for reverse genetics (McCarty and Meeley, 2009). The two major transposon systems used for gene tagging in maize are *Activator/Dissociation* (*Ac/Ds*) and *Mutator* (*Mu*) (Candela and Hake, 2008). These systems consist of an autonomous or master element that encodes a transposase (TPase) and a second non-autonomous or receptor element. The receptor elements are frequently derived from a master element by mutations within the TPase gene. Lacking TPase, non-autonomous elements are stable, unless mobilized by TPase supplied *in trans* by an autonomous element (Kunze et al., 1997). *Ac* is a member of the hAT transposon superfamily (named after the founding members *hobo*, *Ac* and *Tam3*; Calvi et al., 1991) and moves via a cut-and-paste mechanism (Bai et al. 2007), with a preference for transposition to linked sites, making the system ideal for local mutagenesis (Greenblatt, 1984; Dooner and Belachew, 1989; Brutnell and Conrad, 2003). To exploit the *Ac/Ds* system for reverse genetics, *Ds* elements have been distributed throughout the genome to provide potential “launch pads” for mutagenesis of nearby genes (Vollbrecht et al. 2010).

In this study, we identify five genes encoding components of the CDK8 module in maize, present experimentally determined gene structures, and report expression of corresponding transcripts. We performed *Ds* mutagenesis of the gene *ZmMed12a*, identifying two novel insertional alleles, one of which results in a truncation of the ZmMed12a transcript. These insertional mutant alleles will enable determination of the biological roles of the CDK8 module in maize development and stress responses.

## MATERIALS AND METHODS

### Identification of maize CDK8 module genes

Maize CDK8 module genes were identified by BLAST searches using the predicted *Arabidopsis thaliana* protein sequences for HEN3/CDK8 (AT5G63610), CYCC1;1 (At5g48640), CCT/MED12 (At4g00450), and GCT/MED13 (At1g55325) available at TAIR (www.arabidopsis.org). Reciprocal BLAST searches were conducted between all maize and Arabidopsis sequences, to establish that the five maize genes *ZmCDK8*, *ZmCycC*, *ZmMed12a*, *ZmMed12b*, and *ZmMed13* were the only full length CDK8 homologs present in maize.

### Determination of coding sequences for *ZmCDK8*, *ZmCycC*, *ZmMed12a*, *ZmMed12b*, and *ZmMed13*

Multiple mRNA sequences with full-length coding sequences (as well as upstream and downstream untranslated regions) were identified from the NCBI database for both *ZmCDK8* and *ZmCycC.* For *CDK8*, cDNAs for two alternative splice products were identified: EU968864, NM_001157457 and BT018448 correspond to one splice variant, and BT039744 and XR_552425 correspond to the other splice variant. For *CycC*, three independent cDNAs (BT040922, BT033427, and XM008652706) were identified for the one splice variant (shown in Figure 1). Two independent cDNAs (AY105730 and EU972675) represented another *CycC* splice variant with an identical coding sequence but with slight differences in the 3’UTR. A third splice variant was represented by a single cDNA (BT036293); this mRNA has two upstream ORFs, and encodes a truncated CycC protein. For *ZmMed12a*, *ZmMed12b* and *ZmMed13*, partial sequences were obtained from the maize database (maizegdb.org), which were then confirmed and extended by RT-PCR using RNA extracted from seedlings of the B73 inbred line. To confirm the *ZmMed12a*, *ZmMed12b*, and *ZmMed13* gene models, we amplified cDNA products covering the entire predicted coding regions. Given their large expected size, *ZmMed12a*, *ZmMed12b*, and *ZmMed13* cDNAs were amplified in multiple over-lapping fragments. Sequencing of cDNA products was generally consistent with gene models based on genomic sequence analysis, except in the case of *ZmMed13*, where a large intron not present in the maize genome sequence was discovered. Coding sequences were deposited in the NCBI database with the following accession numbers: *ZmMed12a* (KP455660), *ZmMed12b* (KP455661), and *ZmMed13* (KP455662).

**Figure 1.**
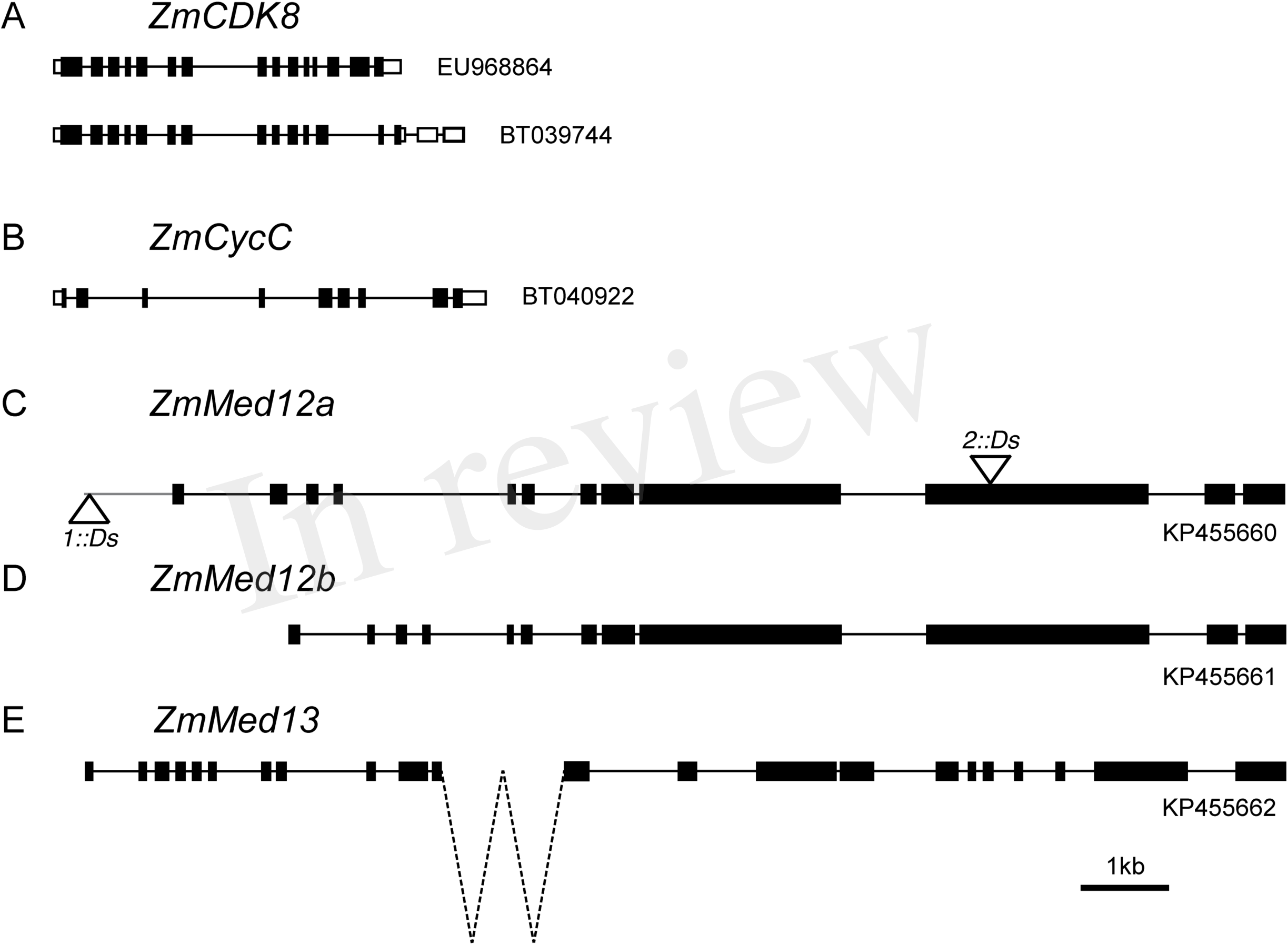
The CDK8 module of maize consists of *CDK8*, *CyclinC*, *Med12a*, *Med12b*, and *Med13*. (A) Exon-intron structure for two different splice products (EU968864 and BT039744) of the *ZmCDK8* gene (GRMZM2G166771). EU968864 encodes a 471AA protein, while BT039744 encodes a 385AA protein, truncated after the CDK8 kinase catalytic domain (cd07842) (B) Exon-intron structure of mRNA sequence BT040922 for the *ZmCycC* gene (gRMZM2G408242), encoding a predicted protein of 257AA. (C) *ZmMed12a* (GRMZM2G114459) encodes a 2193 AA protein (mRNA sequence KP455660). The location of the *Ds* insertions *Zmmed12a-1::Ds* (918bp upstream of ATG) and *Zmmed12a-2::Ds* (exon 10, at bp 4,236 of coding sequence) are indicated. The orientation of each *Ds* insertion is represented by the triangle above the gene (5’-3’), and below (3’-5’). (D) *ZmMed12b* (split gene GRMZM5G828278 / GRMZM5G844080) encodes a 2202 AA predicted protein (mRNA sequence KP455661). (E) *ZmMed13* (split gene GRMZM2G053588 / GRMZM2G153792) encodes a 1892 AA protein (mRNA sequence KP455662). Intron 11 of *ZmMed13* is of unknown size (dotted lines), as it spans a gap in the maize genome sequence. Intron sizes for all genes were determined using corresonding maize genomic sequence. Exons are represented by black boxes, untranslated regions by open boxes, introns by solid black lines, and genomic sequence of ZmMed12a upstream of start codon as solid grey line.

In addition, numerous short genes that are predicted to encode highly truncated ZmMed12 proteins of 199 to 431 residues were identified (Núñez-Ríos, 2012). These short *ZmMed12* genes are predicted to encode the Med12 domain (pfam09497) and many have corresponding expressed sequence tags (EST) (B73 RefGen_v3), which do not cover the entire body of these short genes. Analysis of genomic sequences around these predicted coding sequences did not identify additional *Med12* exons (data not shown), suggesting that these are indeed truncated versions of *ZmMed12*, and not mis-annotated genes with nearby exons that would constitute the middle and C-terminal portions of Med12 proteins.

### Expression profiles of maize CDK8 module genes

Expression data from 22 maize tissues were obtained from http://qteller.com/qteller3/ on August 2014, in the form of Fragments Per Kilobase of transcript per Million (FPKM). In order to look for correlations between pairs of genes across the tissues, the data was log2 transformed (first adding 1, to avoid the logarithm of 0) and normalized using the normalizeQuantiles function from the limma package (Bolstad et al., 2003).

The expression values were selected for the 5 CDK8 module genes: *CDK8* (GRMZM2G166771), *CycC* (GRMZM2G408242), *Med12a* (GRMZM2G114459), *Med12b1* (GRMZM5G828278), *Med12b.2* (GRMZM5G844080), *Med13.1* (GRMZM2G053588), and *Med13.2* (GRMZM2G153792). Since *Med12b.1* and *Med12b.2* as well as *Med13.1* and *Med13.2* are spliced versions of the same gene, the geometric mean was calculated to obtain an averaged estimate of their expression. These data were employed to produce Figure 2A, using the heatmap.2 function from the gplots package (Warns et al., 2015). All pair-wise combinations of the 5 genes across all tissues were plotted using the generic plot function in R (R Core Team, 2015) (Figure S5). The Pearson correlations for all possible pairs of genes were calculated with the cor function, and these data were used as the empirical null to calculate p-values. Correlations for CDK8 module genes were calculated separately. The blob plot in Figure 2B was generated with the corrplot for R.

**Figure 2.**
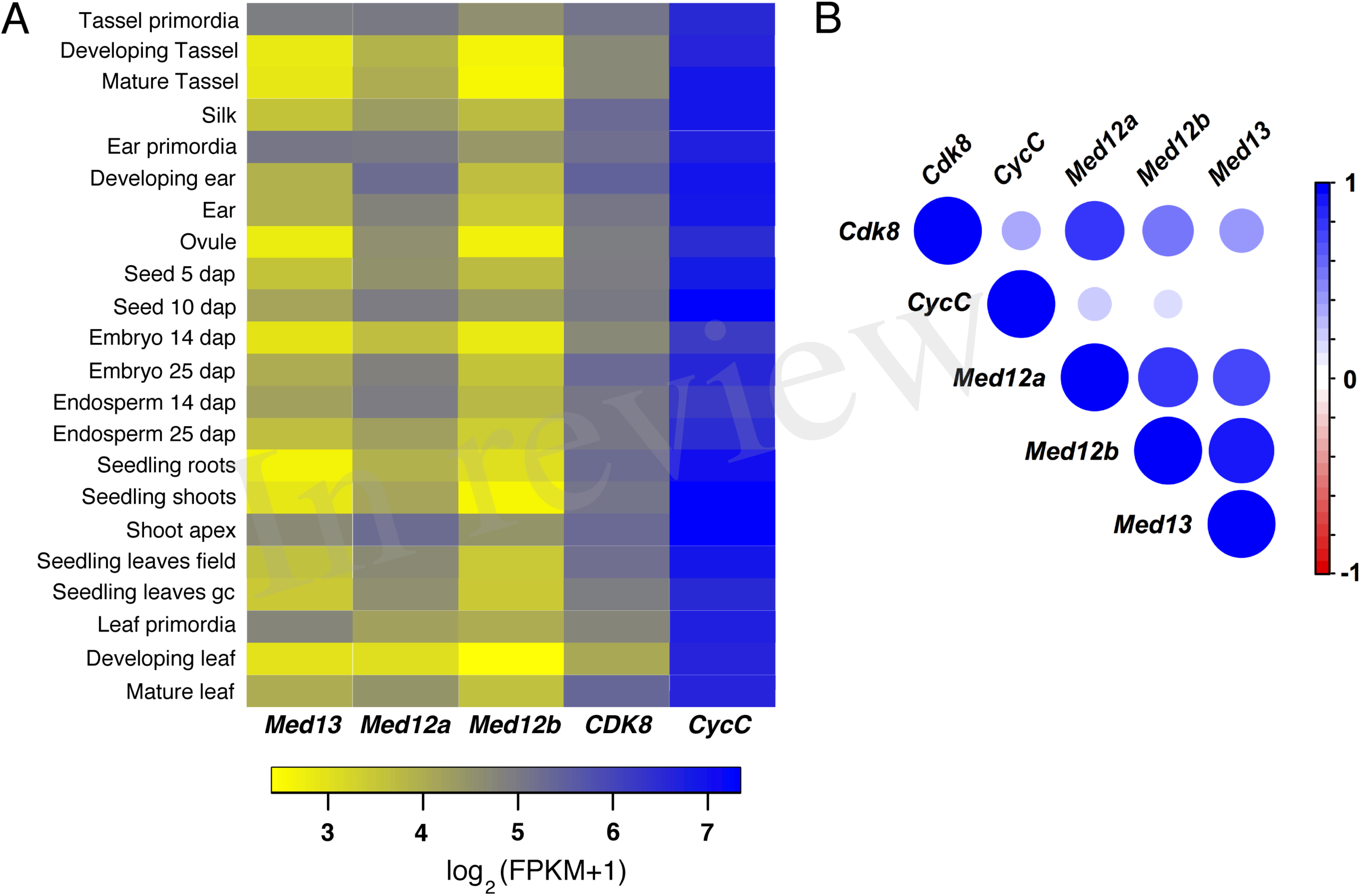
CDK8 module genes are broadly expressed in development. (A) Expression of *Med13*, *Med12a*, *Med12b*, *CDK8* and *CycC* are shown as log2 (FPKM+1) (Fragments Per Kilobase of exon per Million reads mapped). Data are from the following sources: Mature tassel, Developing ear, Ovule, Seed 5 dap, Seed 10 dap, Embryo 25 dap, Endosperm 25 dap, Silk, Developing tassel, Ear, Seedling leaves field, Seedling leaves gc (growth chamber) from Davidson et al. (2011). Developing leaf and Mature leaf from Li et al. (2010). Seedling roots and Seedling shoots from Wang et al. (2009). Embryo 14 dap and Endosperm 14 dap from Waters et al. (2011). Shoot apex, Ear primordia, Tassel primordia and Leaf primordia from Bolduc et al. (2012). (B) Correlation of expression patterns for pairwise combinations of members of CDK8 module. Positive correlations are shown as blue circles, with larger circles and darker blue signifying greater correlations between the two genes. Gene-by-gene comparisons for all tissue samples are shown in Figure S5, from which r values to make this plot were taken.

### Description of maize stocks

All stocks were maintained in the common genetic background of a color-converted W22 inbred line (Dooner & Kermicle, 1971). A stable source of *Ac* transposase was provided by *Ac-immobilized* (*Ac-im*), an *Ac* derivative which has lost 10bp at the 5’ end of the element, preventing excision (Conrad and Brutnell, 2005). Activity of *Ac* transposase was monitored using the mutable *Ds* reporter *r1-sc:m3* that carries a Ds6-like insertion in the *r1* locus that controls anthocyanin production in the aleurone and scutellum tissues (Alleman and Kermicle, 1993): when *Ac* transposase is present, excision of *Ds* from *r1* restores gene function producing colored sectors (Brutnell & Dellaporta, 1994). The donor *Ds* (*dDs*) stock *dDs-B.S07.0835* was generated by isolation of novel transpositions from *r1-sc:m3* as previously described (Vollbrecht et al., 2010). Presence of *dDs-B.S07.0835* was assayed by PCR as previously described (Vollbrecht et al., 2010) using a combination of the *Ds* end primer JSR05 and a primer specific to the genomic site of B.S07.0835 (5’- GACGCACACACGTCAGTATAG-3’). To generate the test-cross population, plants verified as carrying the donor *dDs-B.S07.0835* with *Ac-im* in the genetic background were used as males to pollinate *r1-sc:m3/r1-sc:m3* female plants.

### Seedling screen for transposon insertions in *ZmMed12a*

Testcross progeny were germinated and screened for novel insertions of *Ds* in *ZmMed12a* using a PCR-based strategy. Tissue was collected between 7 and 10 days after planting from pools of 10-18 seedlings using a ≈3mm hole punch, and DNA was isolated following a CTAB-based extraction protocol (Weigel and Glazebrook, 2009). A total of 10 *ZmMed12a* gene-specific primers were designed, covering a region extending from 1.8kb upstream of the translational start to the stop codon. These were used in conjunction with the 5’ and 3’ *Ds*-end primers JSR01 and JGp3, respectively, to amplify DNA adjacent to novel *Ds* insertions in *ZmMed12a* (Table 1). Pools amplifying a product were de-convoluted by screening individuals separately; this second round of PCR used DNA extracted from a different seedling leaf than that sampled for the pool to reduce the chances of recovering somatic transposition events. The PCR products of the second PCR were cleaned (Sambrook and Russell, 2006) and the DNA concentration was adjusted for sequencing by the GENEWIZ Company (South Plainfield, New Jersey, USA). Seedlings carrying putative *med12a* insertional alleles were grown to maturity and propagated by both self-pollination and out-crossing to W22 and B73 inbred lines.

**Table 1:**
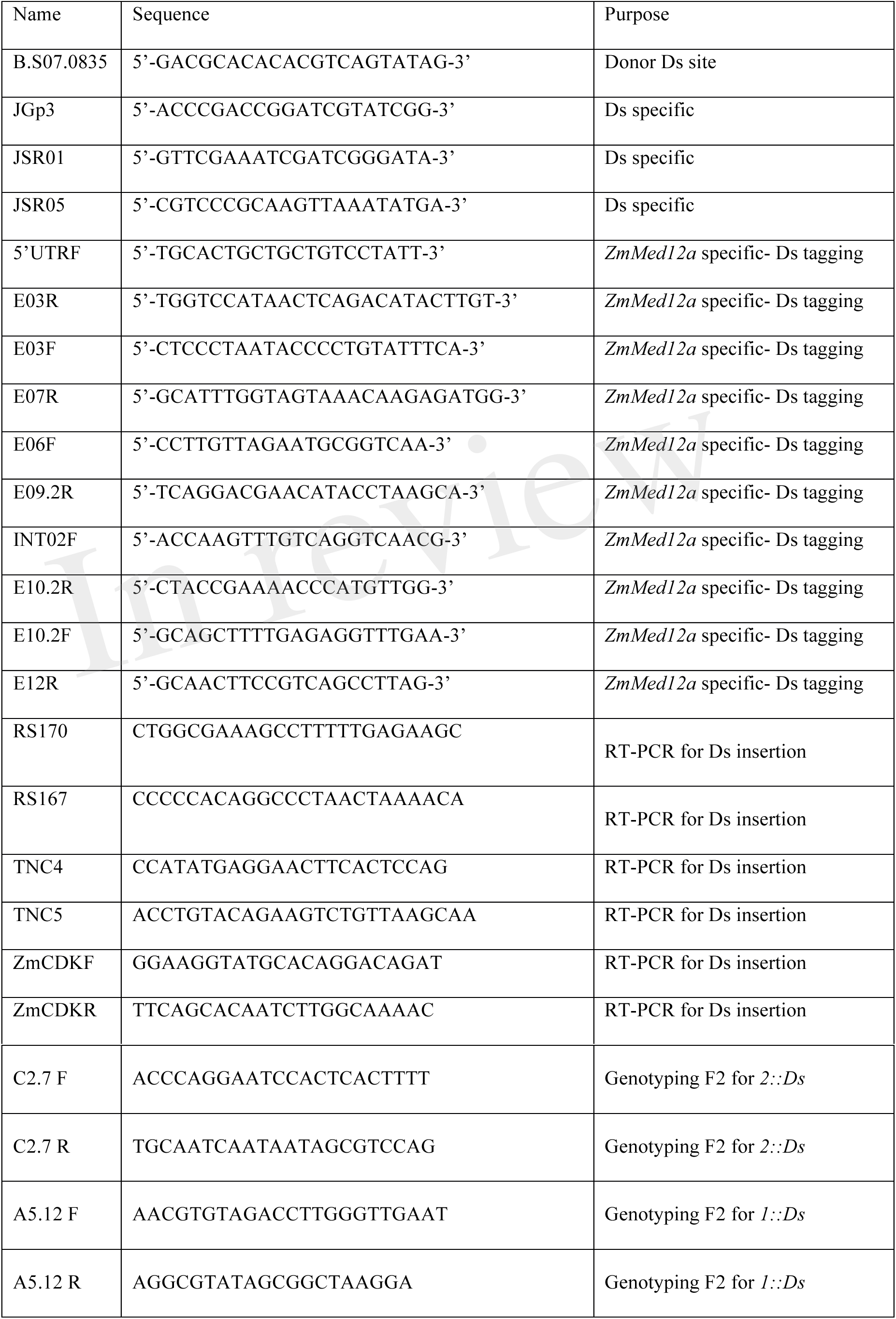
PCR primers used in this study

### RT-PCR analysis of *zmmed12a-1::Ds* and *zmmed12a-2::Ds* alleles

DNA was extracted from 10 day old greenhouse grown seedlings of F2 populations segregating the *1::Ds* and *2::Ds* insertions. Seedlings were genotyped using primers to identify homozygous wild type and homozygous insertion alleles for *1::Ds* (primer pair A5.12F and A5.12R for wild type and A5.12F and JGp3 for *Ds* insertion) and *2::Ds* (primer pair C2.7F and C2.7R for wild type allele and C2.7R and JGp3 for *Ds* insertion). RNA was then extracted Trizol (Invitrogen) for wild type and homozygous insertion alleles. Reverse transcription was performed with SuperScript II (Invitrogen). PCR was performed with the following programs, using Kapa Taq Polymerase (Kapa Biosystems). TNC4-TNC5 primer pair: initial denaturation 95 °C 5′; 10 cycles of 95 °C 30″, 60 °C 30″ (-0.5 °C per cycle), 72 °C 45″; 27 cycles of 95 °C 30″, 55 °C 30″, 72 °C 45″; final extension 72 °C 5′. RS170- RS167 and ZmCDK primer pairs: initial denaturation 95 °C 5′; 30 cycles of 95 °C 30″, 60 °C 30″, 72 °C 1′; 72 °C 10′.

## RESULTS

### The maize genome encodes all four components of the CDK8 module of Mediator

A previous effort to identify Mediator genes from many plant species identified a single maize homolog for all four CDK8 module genes (*CDK8, CYCC, MED12* and *MED13*) (Mathur et al., 2011). In order to conclusively define the number and identity of CDK8 module homologs in maize, we performed BLAST searches to identify all maize gene models (B73 reference genome v3; www.maizesequence.org) whose putative protein products exhibit a high degree of similarity to the entire predicted *Arabidopsis* proteins of the CDK8 module of Mediator: CDK8 (encoded by *HEN3*) (Wang and Chen, 2004); CYCC1;1 or CYCc1;2 (Wang et al., 2004); MED12 (encoded by *CCT/CRP*) (Gillmor et al., 2010; Imura et al., 2012); and MED13 (encoded by *GCT/MAB2*) (Gillmor et al., 2010; Ito et al., 2011) (Table 2). Using the translated experimentally verified coding sequences for all maize CDK8 module genes (see below), all potential orthologous relationships were further validated by reciprocal searching of the *Arabidopsis* genome using maize sequences, and by inspection of the next-best-hit in both *Arabidopsis*-to-maize and maize-to-*Arabidopsis* searches (data not shown).

**Table 2:**
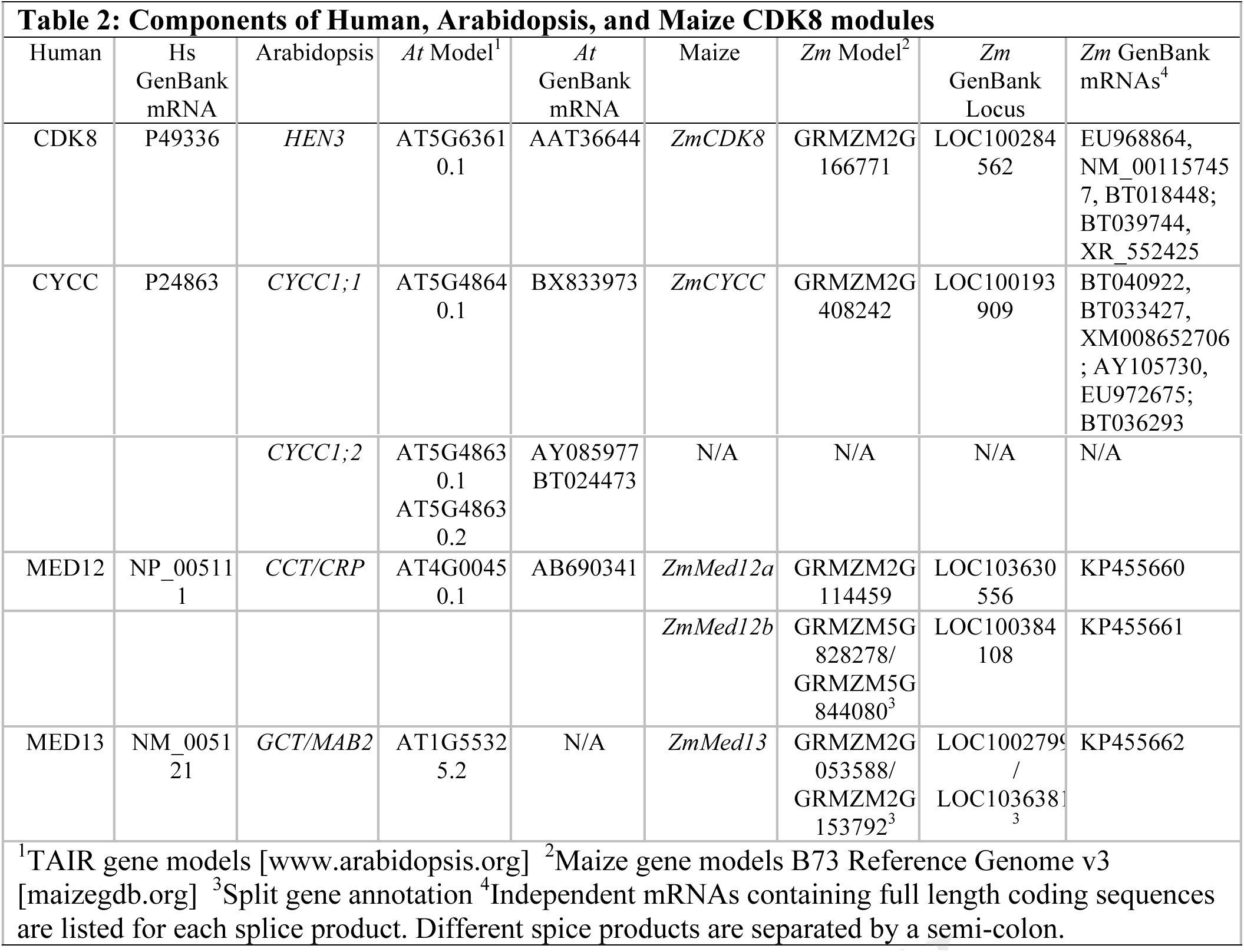
Components of Human, Arabidopsis, and Maize CDK8 modules

| Table 2: Components of Human, Arabidopsis, and Maize CDK8 modules |  |  |  |  |  |  |  |  |
| --- | --- | --- | --- | --- | --- | --- | --- | --- |
| Human | Hs<br>GenBank<br>mRNA | Arabidopsis | *At* Model<sup>1</sup> | *At*<br>GenBank<br>mRNA | Maize | *Zm* Model<sup>2</sup> | *Zm*<br>GenBank<br>Locus | *Zm* GenBank<br>mRNAs<sup>4</sup> |
| CDK8 | P49336 | *HEN3* | AT5G6361<br>0.1 | AAT36644 | *ZmCDK8* | GRMZM2G<br>166771 | LOC100284<br>562 | EU968864,<br>NM_00115745<br>7, BT018448;<br>BT039744,<br>XR_552425 |
| CYCC | P24863 | *CYCC1;1* | AT5G4864<br>0.1 | BX833973 | *ZmCYCC* | GRMZM2G<br>408242 | LOC100193<br>909 | BT040922,<br>BT033427,<br>XM008652706<br>; AY105730,<br>EU972675;<br>BT036293 |
|  |  | *CYCC1*;2 | AT5G4863<br>0.1<br>AT5G4863<br>0.2 | AY085977<br>BT024473 | N/A | N/A | N/A | N/A |
| MED12 | NP_00511<br>1 | *CCT/CRP* | AT4G0045<br>0.1 | AB690341 | *ZmMed12a* | GRMZM2G<br>114459 | LOC103630<br>556 | KP455660 |
|  |  |  |  |  | *ZmMed12b* | GRMZM5G<br>828278/<br>GRMZM5G<br>844080<sup>3</sup> | LOC100384<br>108 | KP455661 |
| MED13 | NM_0051<br>21 | *GCT/MAB2* | AT1G5532<br>5.2 | N/A | *ZmMed13* | GRMZM2G<br>053588/<br>GRMZM2G<br>153792<sup>3</sup> | LOC1002799<br>/<br>LOC1036381<sup>3</sup> | KP455662 |
| <sup>1</sup>TAIR gene models [www.arabidopsis.org] <sup>2</sup>Maize gene models B73 Reference Genome v3 [maizegdb.org] <sup>3</sup>Split gene annotation <sup>4</sup>Independent mRNAs containing full length coding sequences are listed for each splice product. Different splice products are separated by a semi-colon. |  |  |  |  |  |  |  |  |

A single maize gene (GRMZM2G166771) was identified as a potential ortholog of *HEN3/CDK8*, and designated *ZmCDK8*. Two different full-length splice products were identified for this gene (EU968864 and BT039744), predicted to encode a full-length and a truncated maize CDK8 protein (Figure 1A; Figure S1). The full-length ZmCDK8 protein is 471 amino acids (AA), and shows 73% identity with the 470 AA Arabidopsis CDK8 protein, and 43% identity with the 464 AA human CDK8 protein (Figure S1). The smaller ZmCDK8 protein is 385 AA, primarily because of a truncation of the C terminal domain, and shows 75% identity with Arabidopsis CDK8, and 43% identity with human CDK8. This truncation occurs after the CDK8 kinase catalytic domain (cd07842), and is thus unlikely to interfere with the kinase function of the protein (Figure S1).

Although Arabidopsis CYCC is encoded by a tandem-duplicated gene pair (Wang et al., 2004), a single potential maize ortholog of *CYCC* (GRMZM2G408242) was identified, and designated *ZmCycC*. Figure 1B shows the splice product represented by the full-length cDNA clone BT040922 (Figure 1B). The 257AA BT040922 protein is 42% identical to human CycC and 67% identical to Arabidopsis CycC1;1 (Figure S2), and contains the Cyclin domain (cd00043) that is present in human and Arabidopsis CycC (Figure S2).

BLAST searches using the Arabidopsis CCT/MED12 protein identified two putative full- length maize genes (GRMZM2G114459 on chromosome 1, and the split gene GRMZM5G828278 / GRMZM5G844080 on chromosome 9), which were designated *ZmMed12a* and *ZmMed12b*. Partial cDNA sequences were publicly available for *ZmMED12a* and *ZmMed12b;* these sequences, as well as coding sequences predicted by the maize database, were used to experimentally determine mRNA sequences for both genes by RT- PCR. The exon-intron structure of both genes is very similar, with the only differences occurring in the length and position of exons 2, 3 and 4 (Figure 1C&D). These splicing differences lead to several small insertions or deletions in the N-terminal portions of the ZmMed12 proteins, with *ZmMed12a* encoding a protein of 2193AA, and *ZmMed12b* encoding a protein of 2202AA; the two ZmMed12 proteins are 91% identical (Figure S3). ZmMed12a is 19% identical to human Med12, and 46% identical to Arabidopsis MED12; ZmMed12b is 20% identical to human Med12, and 46% identical to Arabidopsis MED12 (Figure S3). The region of highest identity is that comprising the Med12 domain (pfam09497), located at the N-terminus of the Med12 proteins (Figure S3).

A single maize gene was identified corresponding to *GCT/MED13* (split gene GRMZM2G053588 / GRMZM2G153792), and designated *ZmMed13*. Partial cDNA sequences were publicly available for *ZmMed13*; these sequences were used as the basis for RT-PCR experiments to identify full-length mRNA and coding sequences, which demonstrated that *ZmMed13* encodes a protein of 1892 AA, with 20% identity to human Med13, and 49% identity to Arabidopsis MED13 (Figure 1E & Figure S4).

### Maize CDK8 module genes are expressed throughout development

In other organisms where the CDK8 module has been studied, the gene pairs *CDK8* and *CyclinC;* and *Med12* and *Med13*, have similar expression patterns and mutant phenotypes (Yoda et al., 2005; Loncle et al., 2007; Gillmor et al., 2010; Gillmor et al., 2014). In order to determine whether the *CDK8* / *CycC* and *Med12* / *Med13* genes have similar expression patterns in maize, we used publicly available RNA sequence data to quantify CDK8 module gene expression in different tissues and at different developmental stages (see Materials and Methods). As seen in the heatmap in Figure 2A, *CycC* was expressed at much higher levels in all tissues than the other CDK8 module genes, with *CDK8* and *Med12a* the next highest expressed genes, and *Med13* and *Med12b* with the lowest expression levels

In order to more precisely compare tissue-specific expression between the different CDK8 module genes, we made pairwise comparisons for all five genes (Figure 2B & Figure S5). Expression was most highly correlated for *Med13* and *Med12b* (Pearson’s r = 0.93), where the expression ratio between the two genes was close to 1 (compare dotted red line for r, with solid black line representing a 1:1 expression ratio) (Figure 2B & Figure S5). *Med12a* and *Med12b* (r = 0.77); *Med12a* and *Med13* (r = 0.7); and *CDK8* and *Med12a* (r = 0.76) also had high Pearson’s coefficients for pairwise comparisons (Figure 2B & Figure S5). By contrast, *CycC* showed almost no correlation with any of the other CDK8 module genes (Figure 2B & Figure S5). The fact that *CycC* shows little expression correlation with the other CDK8 module genes, and is expressed at higher levels than *CDK8*, and many times higher than *Med13*, *Med12a* and *Med12b*, suggests that *CycC* may play more varied roles in development and physiology than the other CDK8 module genes.

### Maize Med12 is encoded by the duplicated gene pair *ZmMed12a* and *ZmMed12b*

The high degree of similarity between *ZmMed12a* and *ZmMed12b* suggests that they are the result of a recent duplication event (Figure S6). *ZmMed12a* and *ZmMed12b* are located in homologous regions of the genome (1S and 9L, respectively), which derive from a polyploidy event that occurred 5-12 million years ago, sometime after the divergence of maize and sorghum lineages. Although gene loss has reduced the number of genes in present-day maize close to pre-duplication levels, in certain cases both syntenic paralogs have been retained (Schnable et al., 2011). Further inspection revealed a sorghum*Med12* gene (Sb01g050260; *SbMed12*) to be present in a region on Chromosome 1L syntenic to the two maize *ZmMed12* containing regions. Moving up- and downstream from *SbMed12*, micro-synteny was conserved, although, typically, for any given sorghum gene only one candidate ortholog was identified in maize, in either the 1S or 9L region, presumably as the result of gene-loss within paralog pairs following whole genome duplication (Fig. 3).

**Figure 3.**
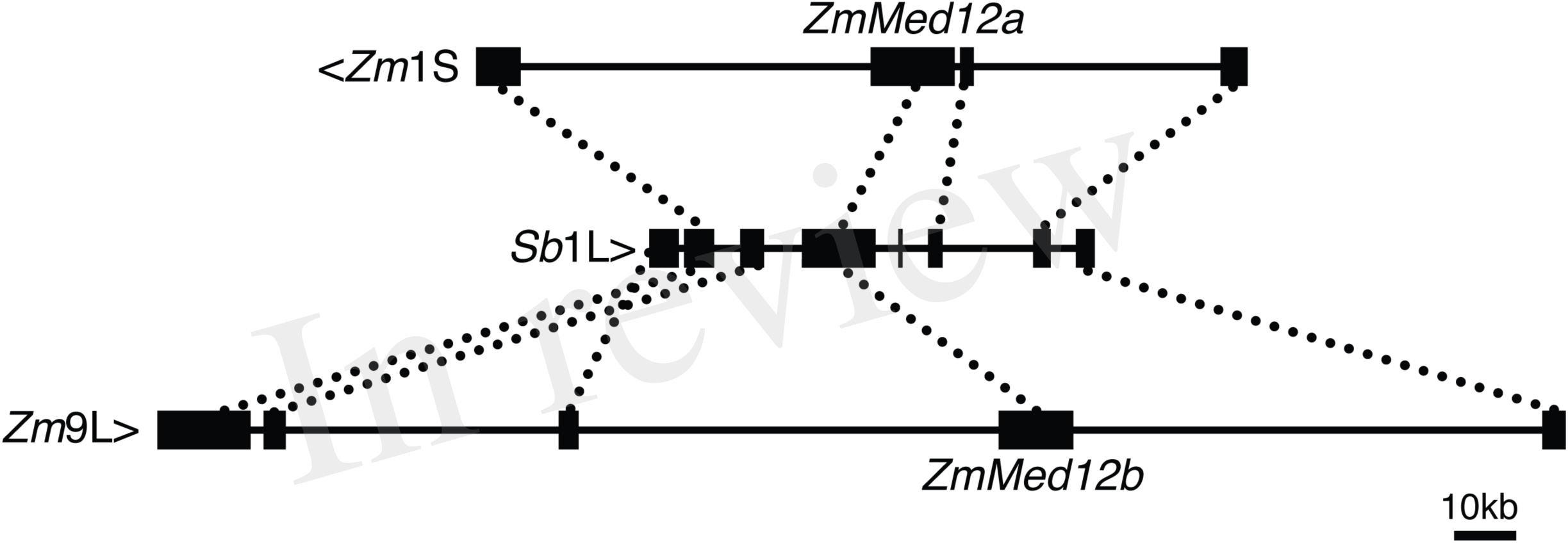
Synteny between maize and sorghum genomic regions surrounding *Med12*. The *Med12* gene is conserved across sorghum and maize syntenic regions. Upper and lower rows: annotated genes in syntenic regions on maize chromosome 1S (*Zm*1S) at ~2Mb (upper row), and maize chromosome 9L (*Zm*9L) at ~155Mb (lower row). Middle row: annotated genes in the region of *SbMed12* at ~73Mb on sorghum chromosome 1L (Sb1L). Orthologous genes are connected by dashed lines. Sb1L and *Zm*9L run left to right, *Zm*1S runs right to left. Genes are shown as black boxes, and the chromosomes are represented by vertical lines. Regions shown to scale, with the right hand position corresponding to the chromosome location mentioned.

### Reverse genetics strategies to target maize CDK8 components

To initiate functional analysis of the maize CDK8 module, we identified publicly available seed stocks carrying *Ac/Ds* or *Mu* family transposons inserted into, or close to, maize CDK8 module encoding genes (Table 3). On the basis of this search, we selected *ZmMed12a* as our first target for reverse genetics: at ˜56kb, the closest potential *Ds* donor was nearer to *ZmMed12a* than to any of the other genes. In addition, the availability of a well-characterized *med12* mutant in *Arabidopsis* provides possibility for comparative study (Gillmor et al., 2010; Imura et al., 2012; Gillmor et al., 2014). Finally, the retention of two *Med12* syntenic paralogs in maize suggests that the roles of *ZmMed12a* and *ZmMed12b* are functionally different, a question which can be addressed by characterization of maize *med12* mutant alleles.

**Table 3:**
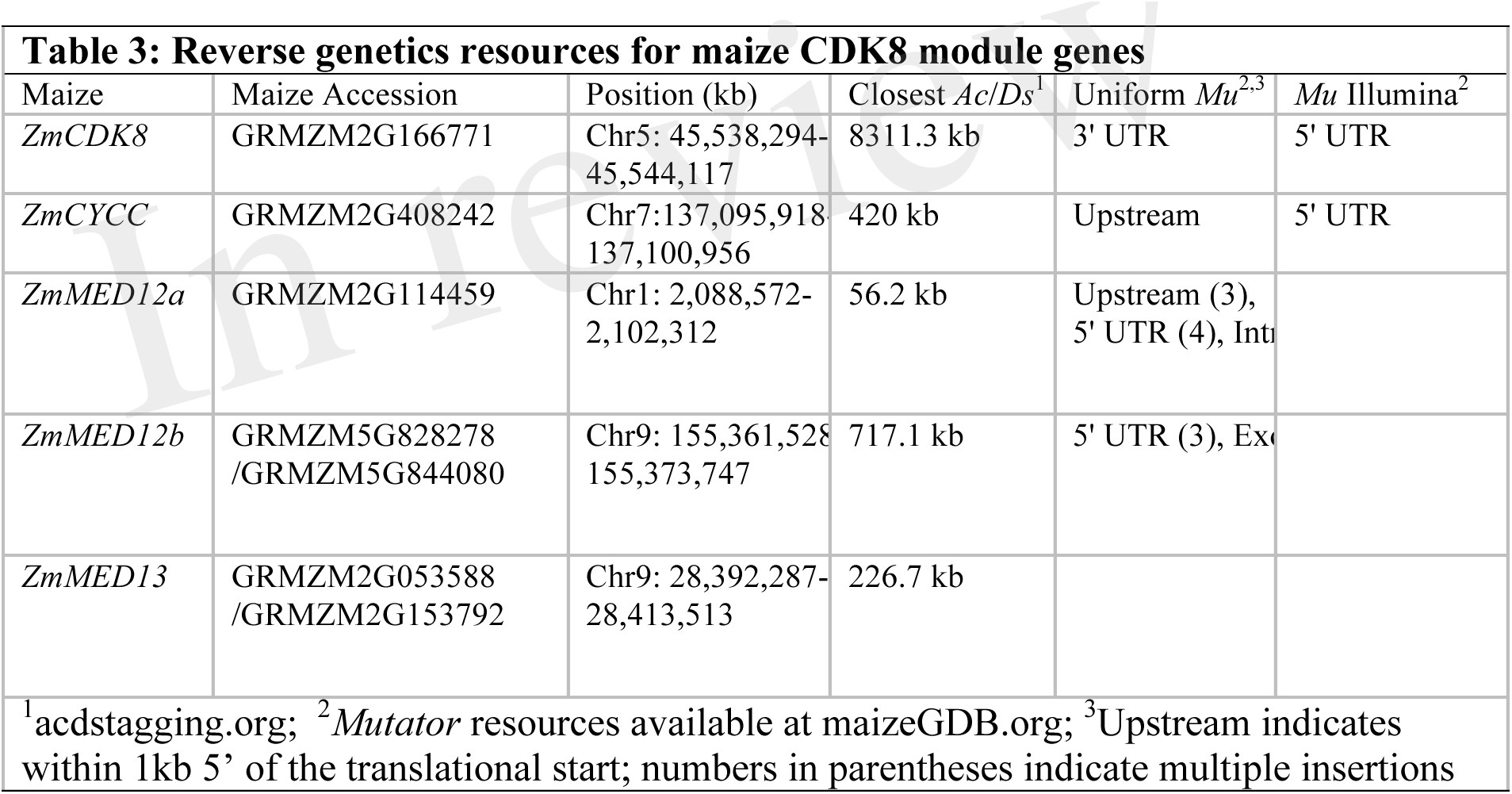
Reverse genetics resources for maize CDK8 module genes

| Maize | Maize Accession | Position (kb) | Closest *Ac/Ds*<sup>1</sup> | Uniform *Mu*<sup>2,3</sup> | *Mu* Illumina<sup>2</sup> |
| --- | --- | --- | --- | --- | --- |
| *ZmCDK8* | GRMZM2G166771 | Chr5: 45,538,294-45,544,117 | 8311.3 kb | 3' UTR | 5' UTR |
| *ZmCYCC* | GRMZM2G408242 | Chr7:137,095,918-137,100,956 | 420 kb | Upstream | 5' UTR |
| *ZmMED12a* | GRMZM2G114459 | Chr1: 2,088,572-2,102,312 | 56.2 kb | Upstream (3), 5' UTR (4), Int |  |
| *ZmMED12b* | GRMZM5G828278 /GRMZM5G844080 | Chr9: 155,361,528-155,373,747 | 717.1 kb | 5' UTR (3), Ex |  |
| *ZmMED13* | GRMZM2G053588 /GRMZM2G153792 | Chr9: 28,392,287-28,413,513 | 226.7 kb |  |  |

<sup>1</sup>acdstagging.org; <sup>2</sup>*Mutator* resources available at maizeGDB.org; <sup>3</sup>Upstream indicates within 1kb 5' of the translational start; numbers in parentheses indicate multiple insertions

### Identification of novel *Ds* insertions into *ZmMed12a*

To use the *Ac/Ds* transposon system to generate mutant alleles of *ZmMed12a*, we first obtained donor *Ds* (*dDs*) stocks carrying the *Ds* element *dDs-B.S07.0835*, located 56.2 kb from *ZmMed12a* (acdstagging.org). The position of the linked *Ds* element was confirmed by PCR assay (see Materials and Methods) (Conrad and Brutnell, 2005). Presence of *Ac-im* in testcross progenitor seed stocks was monitored by somatic excision of a second *Ds* from the *r1-sc:m3* marker locus, resulting in variegated spotting of the kernel aleurone and scutellar tissues (Figure 4A & B). Spotted kernels were planted and seedlings genotyped for the presence of *dDs* using a PCR assay (Materials and Methods). To generate novel germinal insertions into *ZmMed12a*, individuals carrying the *dD* and the *Ac-im* transposase source were used as males to pollinate T43 *(r-sc:m3/r-sc:m3)* females. A test cross population of 59 ears was obtained for the *ZmMed12a* screen (Figure 4A&B).

**Figure 4.**
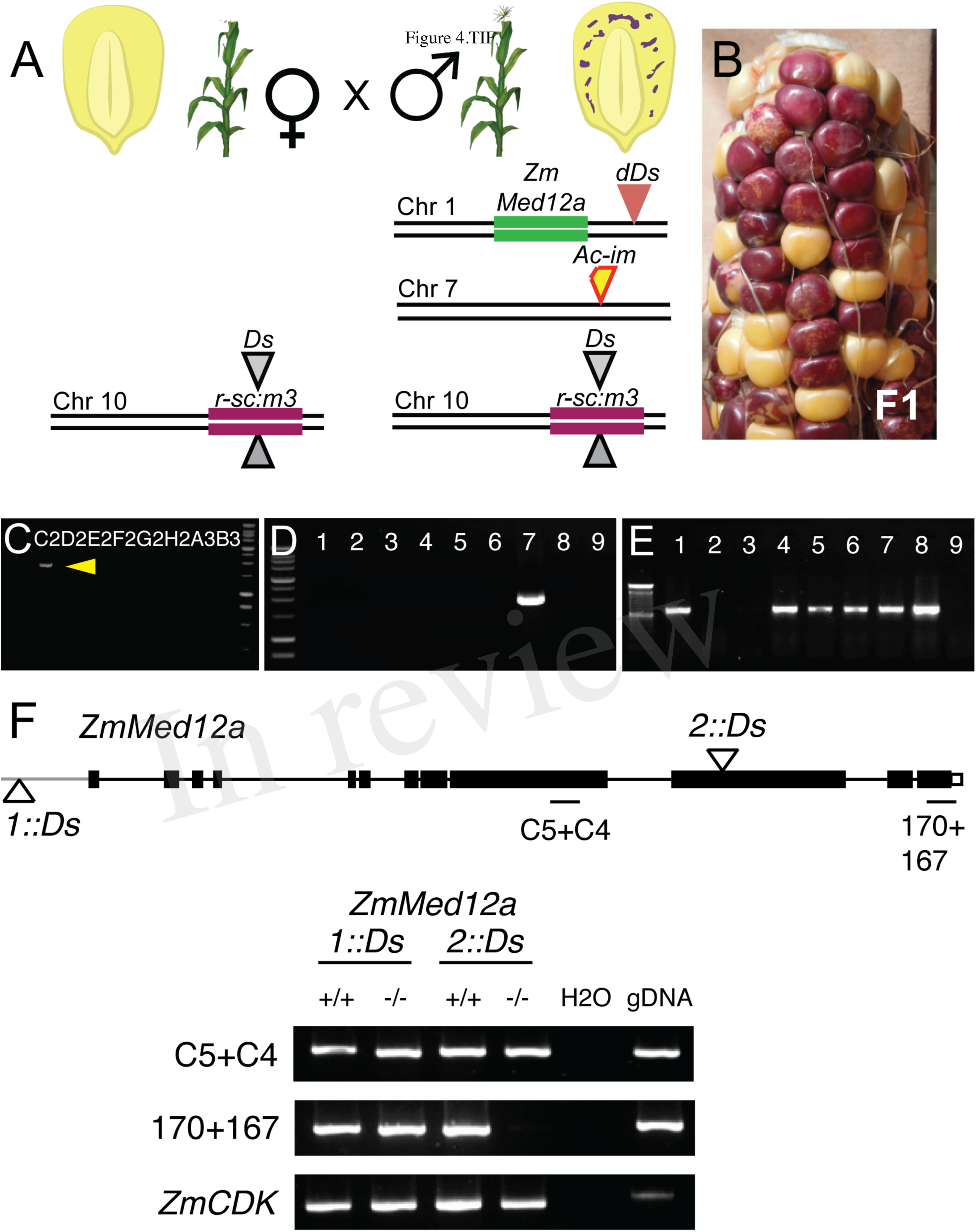
Generation of *Ds* insertional alleles of *ZmMed12a*, and their effect on the *ZmMed12a* transcript. (A) Crossing scheme for generating plants homozygous for the *r- sc:m3* reporter allele, and heterozygous for *Ac-immobilized* (*Ac-im*) and the *Ds* insertion linked to *ZmMed12a*. (B) The presence of *r-sc:m3* allows for selection of spotted F1 kernels, indicating the presence of *Ac-im*, required for remobilizing the *Ds* insertion linked to *ZmMed12a*. (C) Initial pools of 10-18 seedlings were screened by PCR for the presence of a *Ds* element in *ZmMed12a*, using gene specific primer E10.2 and *Ds* specific primer JGp3. A 1.8 kb fragment (yellow arrow) was amplified from pool C2. (D) Individual plants from pool C2 were tested for the presence of the same fragment, which was amplified from plant 7, denominated C2.7 *(Zmmed12a-2::Ds)* (E) This 1.8kb band segregated in nine progeny of the selfed plant C2.7, demonstrating that it is a heritable germinal insertion. (F) RT-PCR analysis of the effect of the *1::Ds* and *2::Ds* insertions on the *ZmMed12a* transcript. The *1::Ds* insertion has no detectable effect on the stability of the *ZmMed12a* transcript, while the *2::Ds* insertion creates a transcript that is truncated after the *Ds* insertion. The location of the primer pairs used to amplify the transcript are indicated in the gene diagram. *ZmCDK* (GRMZM2G149286) was used as a control gene. One representative experiment of three biological replicates is shown.

The test-cross population was screened for *Ds* insertions in *ZmMed12a* using combinations of gene specific and *Ds* specific PCR primers (see Materials and Methods). Pools of 10-18 seedlings were assayed for amplification of putative *Ds*-flanking junction products (see Figure 4C for example for the *Zmmed12a-2::Ds* insertion). Seedlings constituting the pools from which products were amplified were re-screened separately to identify positive individuals (Figure 4D). This second PCR was performed using DNA extracted from a leaf different from that used for the pool PCR to reduce the rate at which we recovered somatic transposition events. We screened a total of 3,049 seedlings and identified two novel insertions into *ZmMed12a: zmmed12a-1::Ds,* located 918bp upstream of the translational start, and *zmmed12a-2::Ds* located in exon 10 (Figure 1C). We performed additional PCR reactions to recover both flanks of the *zmmed12a-1::Ds* and *zmmed12a-2::Ds* insertions. Flanking DNA products were sequenced, confirming the location of the insertions and identifying characteristic 8bp target site duplications. The seedlings carrying the two novel *zmmed12a* insertional alleles were grown to maturity and propagated by both self-pollination and out-crossing. Progeny were germinated and genotyped, confirming the heritability of novel *Ds* insertions (Figure 4E).

### The *2::Ds* insertion results in a truncated *ZmMed12a* transcript

In order to determine the effect of these novel *Ds* insertions on the *ZmMed12a* gene, we performed RT-PCR analysis of plants homozygous for the wild type and *Ds* alleles, using primer pairs that amplify fragments in exon 9 (downstream of the *1::Ds* insertion, and upstrearm of the *2::Ds* insertion), and exon 12 (downstream of both *Ds* insertions) (Figure 4F). Both primer sets amplified fragments from wild type and the *zmmed12a-1::Ds* allele, suggesting that the 1::Ds insertion has no significant effect on the ZmMed12a transcript, a result which is not surprising, since this Ds insertion is upstream of exon 1. In the case of the *zmmed12a-2::Ds* allele, the primer pair in exon 9 produced an amplification product, while the primer pair in exon 12 showed no aplification from homozygous *2::Ds* plants. This result demonstrates that the *2::Ds* insertion causes production of a truncated version of the *ZmMed12a* transcript, likely causing a loss of function of the *ZmMed12a* gene.

## DISCUSSION

In this study we have identified the five genes encoding the CDK8 module of Mediator in maize, determined their coding sequences, characterized their expression in maize tissues during development, and examined the synteny of maize and sorghum in the region of the *Med12* genes. Additionally, we have mutagenized the *ZmMed12a* gene using the *Ac/Ds* transposon system created by Vollbrecht et al. (2010).

In our analysis of CDK8 module genes, we identified two alternative transcripts for *CDK8* (Figure 1). One predicted CDK8 protein is significantly shorter than the other, lacking the C-terminal 86 AA. This truncation seems unlikely to affect enzyme activity *per se*, as the kinase domain is intact (Figure S1). However, the lack of this domain may alter regulation of the kinase activity. Alternatively, the truncation may modify the interaction of CDK8 with CycC, or affect the formation of the four protein CDK8 complex. This complex sterically inhibits the interaction of Core Mediator with RNA pol II, by making direct contact with Core Mediator (Tsai et al., 2013). In the case of CycC, one only one isoform was represented by multiple independent cDNAs. Only single splice products were identified for *Med12a*, *Med12b* and *Med13* (Figure 1). One explanation for this is that there is indeed only one splice product for each gene in maize. It is also possible that the very large size of the mRNAs for these three genes (6-7 kb) makes cloning of multiple splice products difficult, due to technical difficulties in cloning such large cDNAs.

In our analysis of the relative expression of CDK8 module genes, we found *CDK8* and *CycC* to be more highly expressed in all tissues than *Med12a*, *Med12b* or *Med13*. In particular, *CycC* showed the highest expression in all tissues, consistently 3-4 times higher even than *CDK8* (Figure 2). This increased expression of *CycC* is consistent with roles of CycC beyond regulating transcription in tandem with CDK8 (the best known role for CycC) (Allen and Taatjes, 2015). In addition to regulation of transcription, CycC has been shown to promote the G0 to G1 cell cycle transition through phosphorylation of Retinoblastoma, allowing quiescent cells to enter the cell cycle. CycC achieves this through interaction with CDK3, a kinase that is not associated with transcriptional activation, but instead promotes cell cycle entry (Ren and Rollins, 2004). CycC has also been demonstrated to be a haploinsuficient tumor suppressor in mammals, whose loss of function in mice is lethal during embryogenesis (Li et al., 2014). The haploinsuficiency of CycC may require its mRNA or protein levels to be stably maintained, suggesting an explanation for its high levels in all the tissues that we examined (Figure 2 & Figure S6). *Med12a*, *Med12b*, and *Med13* show much lower expression levels, which also vary considerably between different tissues (Figure 2 & Figure S6). The similar expression profiles for *Med12* and *Med13* in maize are consistent with Arabidopsis, where similar expression profiles for these two genes were reported (Gillmor et al., 2010; Ito et al., 2011; Imura et al., 2012; Gillmor et al., 2014). The widely varying expression levels for *Med12* and *Med13* in different tissues are consistent with various roles for these genes in development, both in primordia (where they show the highest expression), as well as in differentiating and more mature tissue.

In Arabidopsis, *MED12* is a single copy gene, with mutant phenotypes in both development and pathogen responses (Gillmor et al., 2010; Imura et al., 2012; Gillmor et al., 2014; Zhu et al., 2014). In maize, however, two *Med12* genes were identified. Sometime after divergence with sorghum, the maize lineage underwent whole genome duplication (Schnable et al., 2011). While in the majority of cases resulting additional gene copies have been lost, for ˜10% of the original gene set syntenic paralog pairs have been retained (Hughes et al., 2015). The genomic location of *ZmMed12a* and *ZmMed12b* is consistent with them representing such a paralog pair. In the region of synteny between maize and sorghum, other genes surrounding *Med12* have been reduced to a single copy, suggesting that the retention of both paralogs of *Med12* in maize may have functional significance. Our isolation of the *2::Ds* insertional allele of *ZmMed12a* will allow us to test the functional importance of this gene. The truncation of the *ZmMed12a* transcript in the *2::Ds* allele makes it very likely that this allele causes a loss of function: a T-DNA insertion in a similar location of the *CCT* (*MED12*) gene of Arabidopsis causes a strong loss of function phenotype, even when some aberrant transcript is produced (Gillmor et al., 2010; Gillmor et al., 2014)

One additional advantage of *Ds* as a mutagen is that novel transpositions occur into linked sites, meaning that the *Ds* insertions in *ZmMed12a* can be remobilized to create further allelic variation in *ZmMed12a*. In addition to mutant alleles that cause a complete loss of function, subsequent *Ds* mutagenesis of *ZmMed12a* may result in hypomorphic alleles that either reduce (but do not eliminate) the function of *ZmMed12a*, or that inactivate specific functional domains of Med12. Alleles that eliminate only certain parts of the Med12 protein could be especially useful in understanding the function of different domains of Med12, currently one of the most interesting, and least explored, aspects of Mediator biology.

## AUTHOR CONTRIBUTIONS

Study designed by TN-R, KA, TPB, SG and RS. Data acquired and/or analyzed by TN-R, KA, ALA-N, DL-S, CM-C, MG-A, SG, RS. Manuscript written by TN-R, SG and RS, and approved by all authors.

## ACKNOWLEDGEMENTS

Thanks to Cei Abreu-Goodger for advice on analysis of gene expression data, and to Jessica Carcaño-Macías for managing seed stocks.

## SUPPLEMENTAL MATERIAL

Supplemental Figures S1-S6.

